# Dynamics of human gut microbiota and short-chain fatty acids in response to dietary interventions with three fermentable fibers

**DOI:** 10.1101/487900

**Authors:** Nielson T. Baxter, Alexander W. Schmidt, Arvind Venkataraman, Kwi S. Kim, Clive Waldron, Thomas M. Schmidt

## Abstract

Production of short-chain fatty acids (SCFAs), especially butyrate, in the gut microbiome is required for optimal health but is frequently limited by the lack of fermentable fiber in the diet. We attempted to increase butyrate production by supplementing the diets of 174 healthy young adults for two weeks with resistant starch from potatoes (RPS), resistant starch from maize (RMS), inulin from chicory root, or an accessible corn starch control. RPS resulted in the greatest increase in total SCFAs, including butyrate. Although the majority of microbiomes responded to RPS with increases in the relative abundance of bifidobacteria, those that responded with an increase in *Ruminococcus bromii* or *Clostridium chartatabidum* were more likely to yield higher butyrate concentrations, especially when their microbiota were replete with populations of the butyrate-producing species *Eubacterium rectale*. RMS and inulin induced different changes in fecal communities, but they did not generate significant increases in fecal butyrate levels.

**IMPORTANCE:** These results reveal that not all fermentable fibers are equally capable of stimulating SCFA production, and they highlight the importance of the composition of an individual’s microbiota in determining whether or not they respond to a specific dietary supplement. In particular, *R. bromii or C. chartatabidum* may be required for enhanced butyrate production in response to RS. Bifidobacteria, though proficient at degrading RS and inulin, may not contribute to the butyrogenic effect of those fermentable fibers in the short term.

## INTRODUCTION

Short-chain fatty acids (SCFAs) are major end products of bacterial fermentation in the human colon and are known to have wide-ranging impacts on host physiology. Butyrate in particular is important for maintaining health via regulation of the immune system (1), maintenance of the epithelial barrier (2, 3), and promoting satiety following meals (4). It may be protective against several diseases, including colorectal cancer (5), inflammatory bowel disease (6), graft-versus-host disease (7), diabetes (8), and obesity (8, 9). Therefore, stimulating butyrate production by the colonic microbiome could be useful for sustaining health and treating diseases.

One strategy for stimulating butyrate production is to supplement the diet with carbohydrates that are resistant to degradation by human enzymes but can be metabolized by select bacteria in the colon. We previously demonstrated that one such resistant starch (RS) prepared from potatoes (RPS) could increase average fecal butyrate in healthy, young adults (10). Others have reported increased butyrate in response to inulin in humans and resistant starch from maize (RMS) in mice (11, 12). A critical challenge to these potential therapies is the variable responses between individuals, likely influenced by differences in the composition of their gut microbiota. To capture this variability large numbers of subjects are required. We analyzed samples from 174 university students who consented to participate in and then successfully completed this short-term interventional study. This young cohort provided a wide diversity of gut communities without the additional complications of chronic health conditions such as obesity, Type 2 diabetes or cardiovascular disease that are related to altered microbiome structure and function (13, 14). We did not ask the participants to make changes to their diets other than taking the supplements provided, even though we recognize different diets also have a profound impact on microbiome structure and function. Our objective was to determine how these different resistant polysaccharides affected the concentrations of SCFAs when added to “normal” diets, not just under one dietary regimen. With this number of participants we were able to evaluate three different resistant polysaccharides and an amylase-sensitive polysaccharide as a negative control.

Understanding the butyrogenic effect of these supplements and specific gut bacteria is important for designing more broadly effective therapies and predicting which individuals are likely to benefit from them. More generally, defining metabolic interactions among gut microbes enhances our understanding of the assembly, maintenance and outputs from the gut microbiome.

Identifying butyrogenic configurations of the microbiome is challenging because several different bacteria (or combinations of bacteria) may be involved in the multi-step process. Many bacteria in the colon are involved in the degradation of dietary fiber, the complex mixture of plant polysaccharides that is not susceptible to host enzymes (15, 16). However, many of these bacteria are specialists, attacking specific bonds in specific types of polymers (17–19). Only a limited number of gut bacteria may be able to degrade any given resistant polysaccharide that is used as a dietary supplement. Primary degraders depolymerize specific polysaccharides to mono-, di- and oligo-saccharides that they can take-up and ferment themselves to acidic end products such as acetate or lactate (20). Their selective growth on the dietary supplement should result in a higher relative abundance in fecal communities. However, most resistant starch degraders are not among known butyrate producers (19, 21). Thus, for these supplements to stimulate butyrate production, the activities of additional organisms would be required. These secondary fermenters capture degradation and fermentation products from primary degraders and metabolize them into new molecules including butyrate (Fig. 1). However, if primary degraders use the supplements efficiently, only a fraction of the carbon and energy they contain may become available to the secondary fermenters. Therefore, increases in the relative abundance of butyrate producers may be more difficult to detect, but their metabolic activity could still be evidenced by an increase in fecal butyrate.

**Figure 1.**
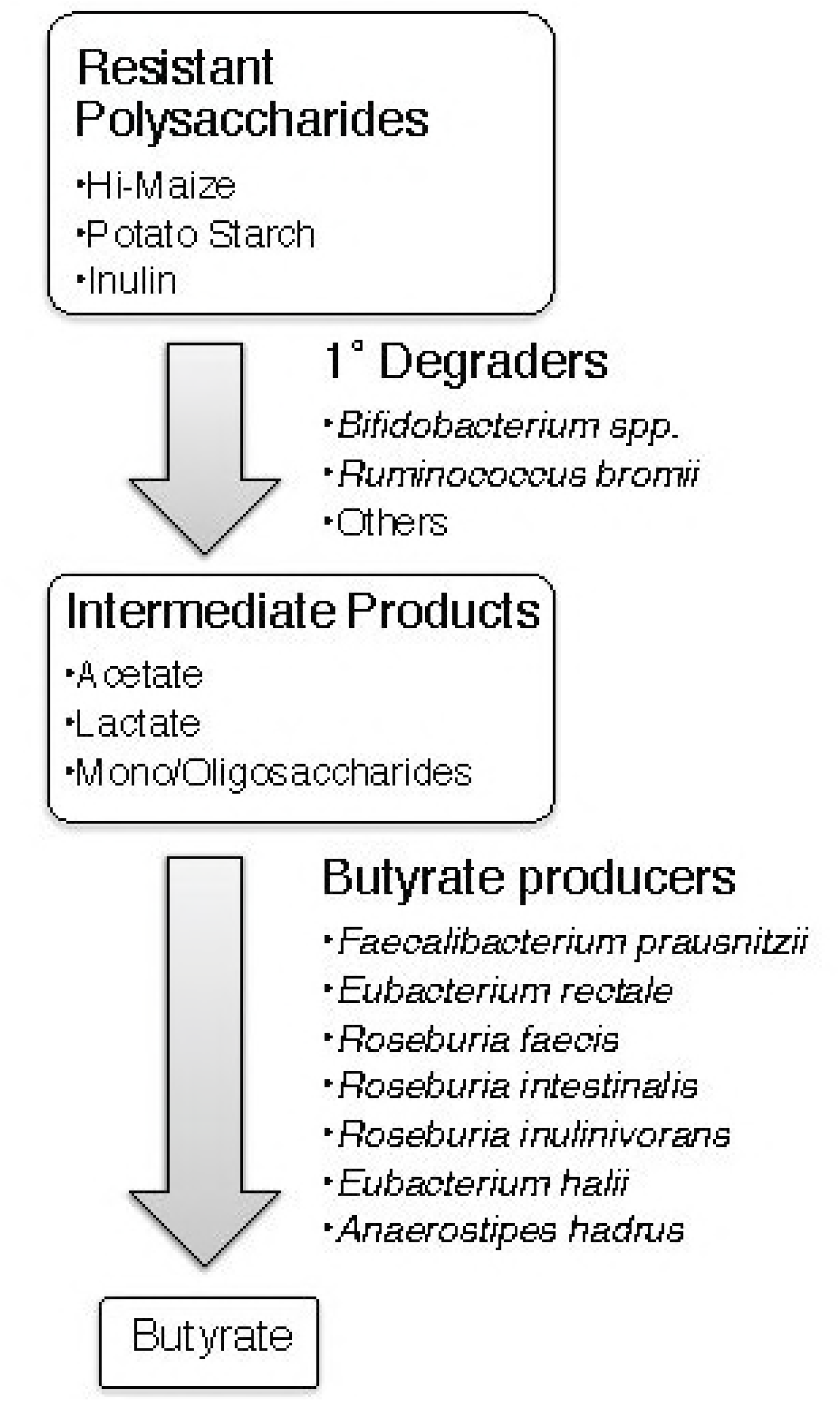
Proposed model of metabolites and microbes that catalyze the flow of carbon from resistant polysaccharides to butyrate. There are cultivated strains from the gut microbiome that possess the metabolic activities proposed for the species listed.

Candidates for performing one or more of the steps in Fig. 1 *in vivo* have already been identified by their metabolic capabilities *in vitro*. For example, *Ruminococcus bromii* and *Bifidobacterium adolescentis* have been shown to degrade resistant starches (22). There have been claims that other species degrade resistant starches, but they are less compelling because they either involved starch preparations that could include sensitive as well as resistant fractions (23) or because the evidence was indirect (e.g. binding to starch granules (24)) or the presence of DNA sequences encoding amylase-like GH13 domains (25). The ability to degrade inulin *in vitro* has been demonstrated for some species of *Bifidobacterium* (though not all strains) (26) and by several members of the *Lachnospiraceae* and *Ruminococcaceae* families in the phylum Firmicutes (27). These two families include most of the known butyrate producers (21). The most abundant of them in the human gut are *Eubacterium rectale* and *Faecalibacterium prausnitzii*, both of which are capable of degrading inulin and producing butyrate from it (27). *E. rectale* has been shown incapable of degrading resistant starches unless they are heat-treated to denature some of the crystal structure (22). Such rigorous tests have not been reported to our knowledge for *F. prausnitzii. In vitro* studies have also demonstrated that combinations of primary degraders and secondary bacteria can produce butyrate from resistant polysaccharides by cross feeding. For example, *Eubacterium rectale* can grow on degradation products of RS released by *R. bromii*, and several species of *Roseburia* and *F. prausnitzii* have been cross-fed by *Bifidobacterium* spp. (20, 22, 28, 29). Bifidobacteria also promote butyrate production by another select group of bacteria because they produce both acetate and lactate via a unique fermentation pathway known as the Bifid shunt (30). This combination of end products can be converted into butyrate by *Eubacterium hallii, Anaerostipes caccae* or *Anaerostipes hadrus* (31, 32).

These known degraders and butyrate producers were targeted for evaluation in this dietary intervention. But we also analyzed the entire community of fecal bacteria to identify any organisms that had not previously been associated with the metabolism of these fermentable fibers. We attempted to address four major issues:

1. Do the three resistant polysaccharides stimulate butyrate production in this population of healthy, young individuals? If so, do they have similar impacts on butyrate production?
2. Which gut bacteria respond to these dietary additions by increasing in relative abundance? Can we identify any species that were unexpectedly affected? Are the same bacteria affected by all three supplements?
3. Can we find any evidence of selectivity, either in the substrates used by primary degraders or in the butyrate producers they cross feed?
4. Do changes in the relative abundance of primary degraders and butyrate producers explain differences in individuals’ butyrate concentrations?

## RESULTS

### Effects on Short Chain Fatty Acids

We first examined the impact of each supplement on the concentration of SCFAs in the feces. Both RPS and inulin significantly increased total SCFA concentrations, by 32% and 12% respectively (both p<0.001). Supplementation with RPS increased butyrate concentrations by an average of 29% (p<0.001) and acetate by an average of 21% (p=0.0012, Table 1). However, the response was highly variable between individuals: the median concentration of butyrate increased in 63% of individuals and was either unchanged or decreased in the remaining 37%. Although total SCFA concentrations increased with inulin supplementation, there were no statistically significant changes in individual SCFAs. Nor were there significant changes in the concentration of any of the SCFAs in the groups whose diet was supplemented with either RMS or accessible starch (Table 1). Furthermore, there were no significant differences in SCFA concentrations between the control group that consumed 20 g of accessible starch compared to the group that consumed 40 g of accessible starch.

**Table 1.** The concentrations of fecal SCFAs (in mmoles/kg) before and during dietary supplements. All p-values are based on repeated measures ANOVA.

| Substrate | Butyrate |  |  |  | Acetate |  |  |  | Propionate |  |  |  |
| --- | --- | --- | --- | --- | --- | --- | --- | --- | --- | --- | --- | --- |
|  | before<br>(mean ± SD) | during<br>(mean ± SD) | Change | p-value | before<br>(mean ± SD) | during<br>(mean ± SD) | Change | p-value | before<br>(mean ± SD) | during<br>(mean ± SD) | Change | p-value |
| Accessible<br>(n=39) | 13 ± 6.1 | 15 ± 8.3 | +13% | 0.18 | 41 ± 17 | 41 ± 16 | 0% | 0.89 | 9.9 ± 6.0 | 9.3 ± 6.5 | -6% | 0.47 |
| Hi-Maize<br>(n=43) | 9.3 ± 4.1 | 9.7 ± 5.6 | +5% | 0.81 | 37 ± 17 | 33 ± 15 | -10% | 0.20 | 12 ± 15 | 12 ± 13 | +0.3% | 0.81 |
| Potato<br>(n=43) | 13 ± 6.0 | 16 ± 7.5 | +29% | **<0.001** | 48 ± 22 | 58 ± 26 | +21% | **0.0012** | 10 ± 7.7 | 8.6 ± 5.3 | -16% | 0.39 |
| Inulin<br>(n=49) | 11 ± 6.0 | 13 ± 7.0 | +17% | 0.14 | 38 ± 18 | 41 ± 20 | +8% | 0.077 | 11 ± 10 | 13 ± 15 | +27% | 0.31 |

### Effects on Bacterial Communities

We characterized changes to the gut microbiota using 16S rRNA gene sequencing. We chose not to cluster sequences into operational taxonomic units (OTUs) after discovering that several taxa of interest, with very different responses to the dietary supplements, would be clustered into a single OTU even at 99% identity. For example, *Bifidobacterium longum* and *B. faecale* have varying abilities to degrade RS, but the V4 region of the 16S rRNA gene in these species is 99.6% identical. Combining sequences corresponding to these species masks a biological pattern that is readily apparent when considering the unique sequences. Unlike OTUs that are calculated *de novo* with each new data set, unique sequences also have the benefit of being directly comparable across datasets. Nevertheless, the small size of the V4 region of *Bifidobacterium* makes it impossible to resolve all species within this genus. *B. adolescentis, B. faecale* and *B. stercoris* have identical V4 regions, as do *B. longum* and *B. breve*. A third group, *B. catenulatum, B. pseudocatenulatum*, and *B. kashiwanohense*, also share identical V4 regions. For all other species of interest a single sequence was identified that was specific to each species. To avoid analysis of spurious sequences, we limited our analysis to the 500 most abundant unique sequences, which accounted for 71% of the approximately 70 million curated sequencing reads. Using this approach we determined that both the RPS and inulin significantly altered the overall structure of the community (PERMANOVA, p=0.001 and p=0.002, respectively), while the accessible starch and RMS did not (p=1.0 and p=0.65, respectively). None of the supplements significantly changed the alpha diversity, as measured by the inverse Simpson index (p>0.05).

### The Most Affected Bacterial Populations

The sequences that changed the most were identified by the ratio of their relative abundance during supplementation to their relative abundance before (Fig. 2). Most of the sequences that significantly increased in relative abundance were from species already known to degrade resistant polysaccharides. RPS increased the relative abudance of *B. faecale/adolescentis/stercoris* sequences 6.5-fold (p<0.001), but there were no significant changes in any of the other sequences classified in the genus *Bifidobacterium* (Fig. 2). RMS resulted in a 2.5-fold increase in the relative abundance of sequences classified as *R. bromii* (p<0.001), but no significant changes in any of those classified as *Bifidobacterium* (Fig. 2, Supplemental Table 2). Inulin significantly increased the relative abundance of each of the four most abundant sequences assigned to a *Bifidobacterium* species (Fig. 2, all p<0.05) and sequences classified as *Anaerostipes hadrus* (Fig. 2, Supplemental Table 3). No bacterial populations significantly changed in response to the accessible starch supplement (Fig. 2, Supplemental Table 4).

**Figure 2.**
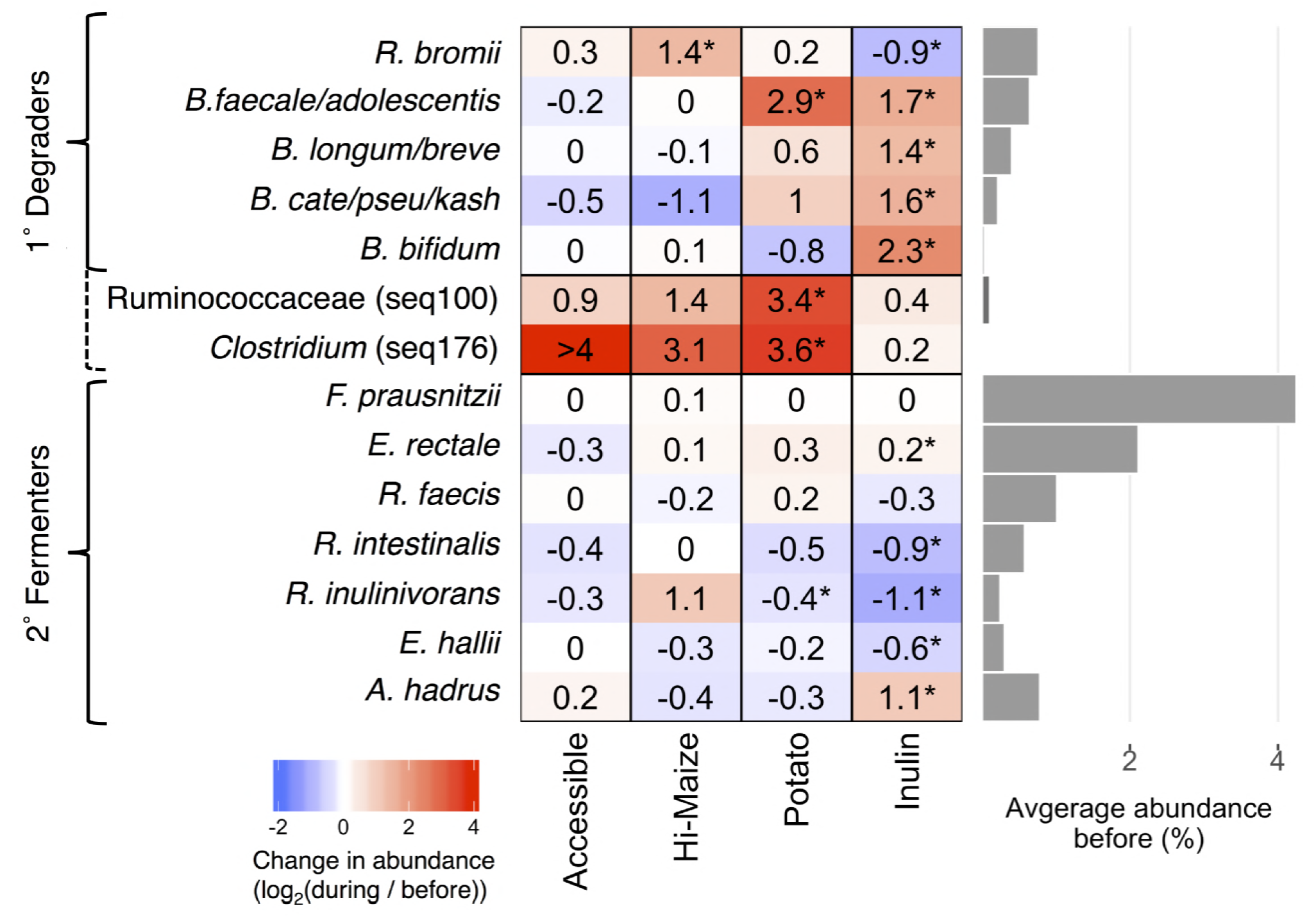
Average fold changes in the relative abundance of sequences representing selected primary degraders of resistant polysaccharides and secondary butyrate fermenters in response to dietary supplements (* p<0.05, paired Wilcoxon test). Seq100 represents an unknown species in the family Ruminococcaceae while seq176 is 98.8% identical to *Clostridium chartatabidum*. Both are inferred to be primary degraders (dashed bracket) based on the dynamics of their response to dietary supplements. Bar plot to the right shows the average relative abundance of each species prior to fiber supplementation.

We also noted increases in sequences that were not significant for the population as a whole, but which were dramatic within a subset of individuals (Supplementary Table 1). RPS increased the relative abundance of *R. bromii* sequences in a subset of individuals (Fig. 3A), but not for the group as whole (p=0.72). One of the most striking increases in a smaller subset was in Seq100, classified as a member of the *Clostridium* cluster IV within the family *Ruminococcaceae. Clostridium leptum* is the closest cultured relative, but their 16S rRNA V4 regions are only 95% identical. This *Clostridium leptum*-related sequence was detected in 11 of the 50 individuals who consumed RPS, increasing by an average of 10-fold and exceeding 10% relative abundance in several individuals (Fig. 3A). Another unanticipated increase was in Seq176 whose V4 region is 98.8% identical to *Clostridium chartatabidum*, a rumen isolate shown to degrade a variety of dietary fibers (33, 34). It was rarely detected before supplementation, but its relative abundance increased up to 4% relative abundance in 11 of the individuals consuming RPS (Fig. 3A). Although a large fold change for Seq176 was observed in the accessible starch and Hi-Maize groups (Fig 3A), it was limited to one individual in each group. Seq176 reached only 0.03% and 0.14% relative abundance in those two individuals after starting below the limit of detection.

**Figure 3.**
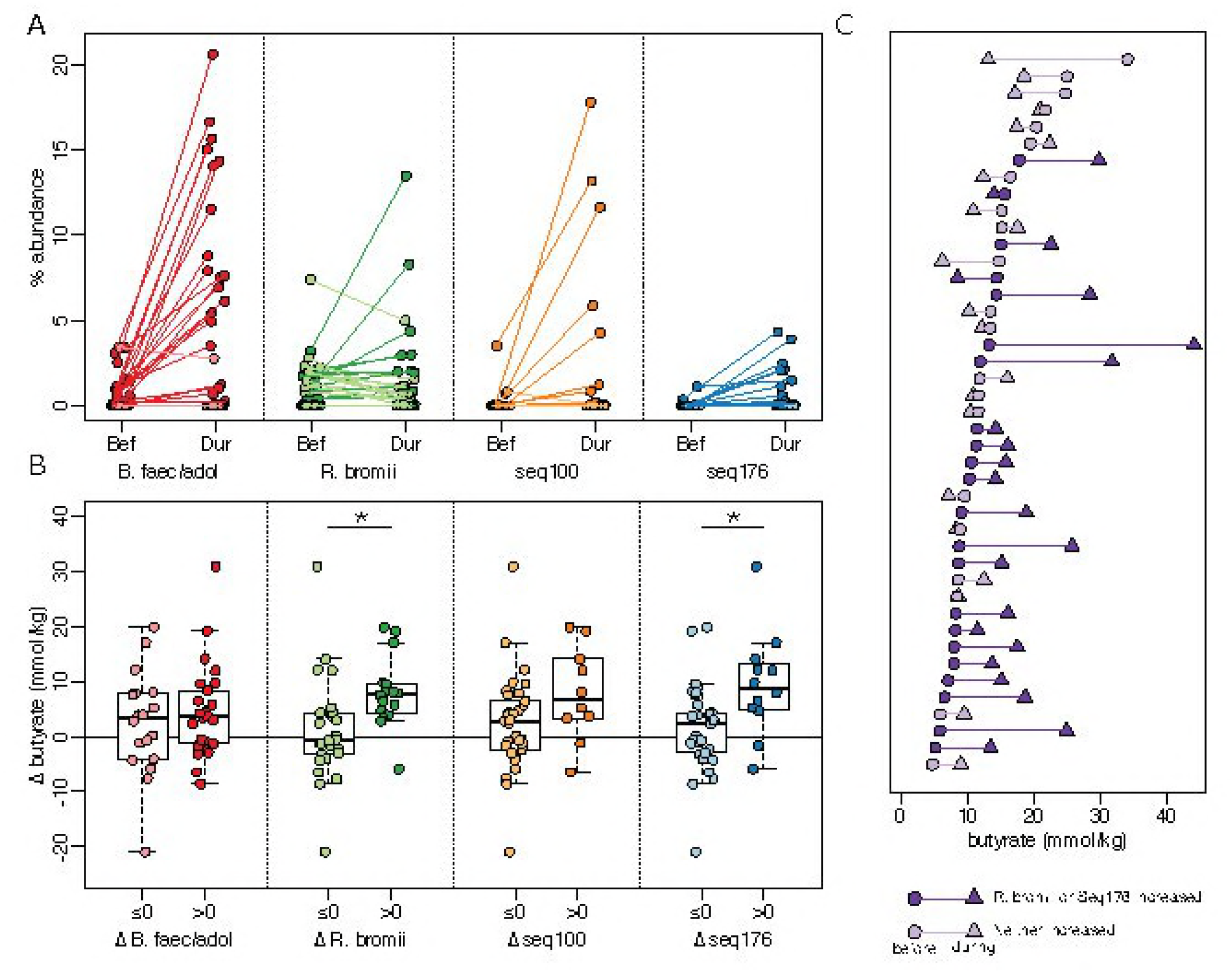
Associations between primary degraders and changes in fecal butyrate concentrations in response to dietary supplementation with resistant potato starch (RPS). For all panels, darker shades indicate an increase in abundance or concentration and lighter shades indicate a decrease or no change. (A) Average relative abundance of putative primary degraders in each individual before (Bef) and during (Dur) RPS supplementation. (B) Change in fecal butyrate in individuals grouped on whether a primary degrader increased (Δ > 0) or did not increase (Δ ≤ 0) in relative abundance in response to RPS supplementation (* p<0.05, t-test). (C) Butyrate concentrations for each individual before (circles) and during (triangles) RPS supplementation. Subjects are sorted by initial butyrate concentration.

### Associations between Butyrate Changes and Community Changes

In individuals consuming RPS, increases in the relative abundance of sequences attributed to either *R. bromii* or *C. charatabidum* were each associated with an increase in butyrate (p=0.025 and p=0.0024, Fig. 3B). On average, there was a 9.1 mmol/kg increase in fecal butyrate when either *R. bromii* or *C. charatabidum* sequences increased in relative abundance in response to RPS. There was a decrease of 2.1 mmol/kg in individuals in whom neither sequence increased (Fig. 3C). Furthermore, the presence or absence *R. bromii* prior to RPS supplementation was indicative of whether an individual would have higher fecal butyrate in response to RPS. Of the 29 individuals with detectable *R. bromii* at baseline, 22 (76%) had higher fecal butyrate during RPS supplementation, compared to 5 (36%) of the 14 individuals without detectable *R. bromii*. The baseline abundance of *C. charatabidum* was not associated with a butyrate response because it was below the limit of detection (<0.008% abundance) in all but five individuals prior to supplementation. With greater sequencing depth or a more sensitive assay it may be possible to use the presence of these two organisms to predict whether an individual will respond to RPS with increased butyrate. Increases in either *B. faecalis/adolescentis/stercoris* or *Clostridium leptum-like* Seq100 were not associated with an increase in butyrate (Fig. 3B).

### Partnerships Converting Polysaccharides to Butyrate

Increasing *R. bromii* was associated with higher butyrate levels, but butyrate is not a major end product of *R. bromii* metabolism (35). Therefore, an increase in *R. bromii* is not sufficient in itself to explain the association with higher butyrate concentrations. Based on our working model (Fig. 1), we expected the increase in primary degraders to lead to an increase in the abundance of butyrate producers (though to a lesser extent because the primary degraders were presumed to extract most of the nutritional value of the supplements for themselves). To test this expectation, we correlated the change in abundance of sequences associated with each of the putative primary degraders with changes in the abundance of the most common known butyrate producers in our cohort (Fig. 4). Consistent with our model, change in *R. bromii* sequences was positively correlated with the change in abundance of *E. rectale* sequences (Fig 4). This observation is also consistent with previous reports that populations of *R. bromii* and *E. rectale* are associated with each other, both physically and metabolically (22, 24). The relative abundance of *E. rectale* was also correlated with the concentration of butyrate in the RPS group (Spearman rho=0.42, p<0.001, Fig. 5), which would explain the higher butyrate in individuals where *R. bromii* increased.

**Figure 4.**
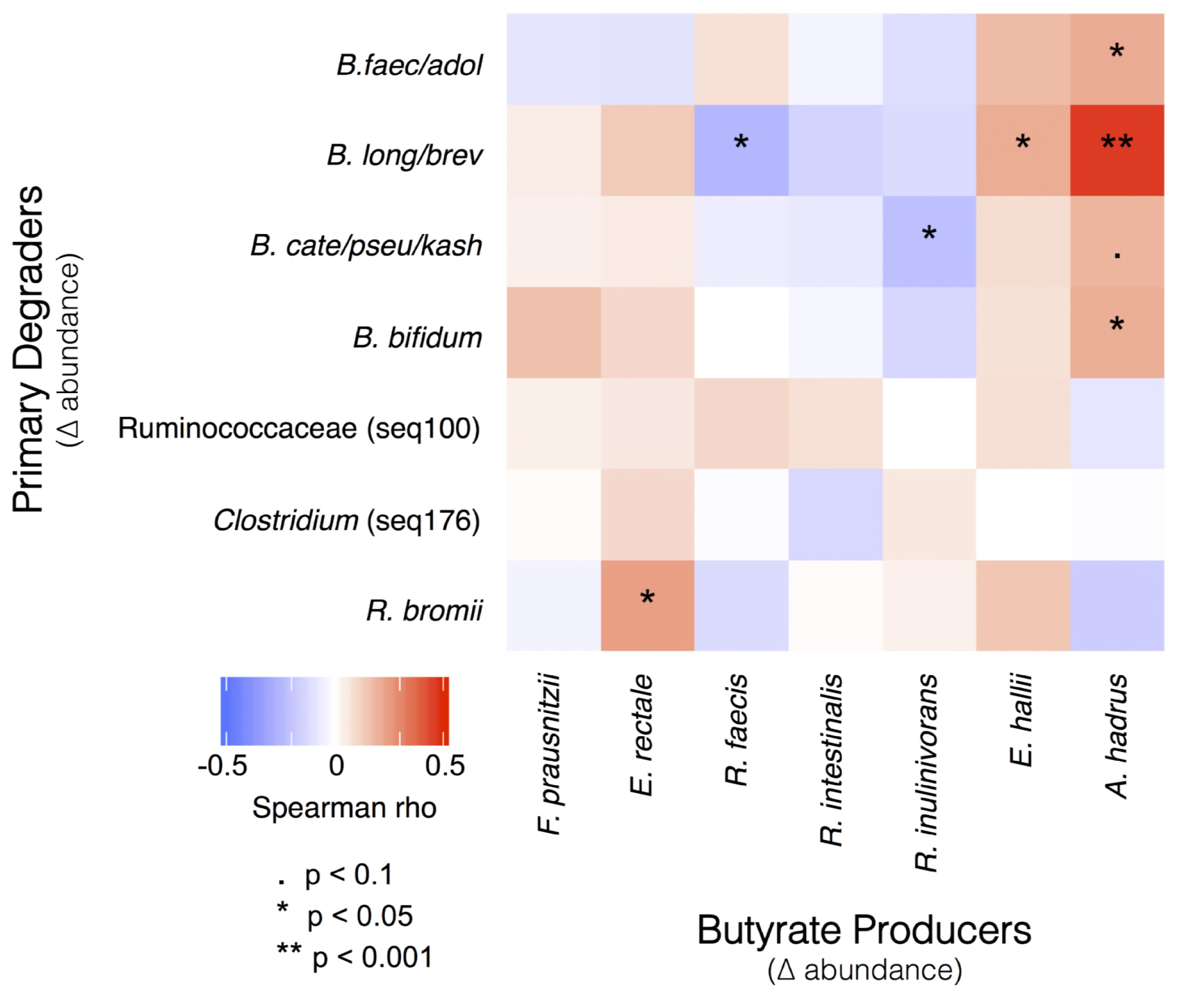
Pairs of microbes that consistently responded in concert either positively (red) or negatively (blue) to dietary supplementation. Correlations between changes in the abundance of primary degraders and butyrate producers were calculated using the combined dataset that includes responses to all supplements.

## DISCUSSION

We tested the effects of fermentable fiber supplements on the structure and function of bacterial communities in the human colon. The three supplements we studied include fractions that are resistant to degradation by host enzymes and therefore pass through the small intestine to the colon. There they can be metabolized by specialized bacteria (which distinguishes them from “bulking fibers” that promote regularity but pass through both small and large intestines without degradation). Two of the supplements we tested were Type 2 Resistant Starches: unmodified starches extracted from potato tubers (RPS) or hi-amylose maize seeds (RMS). However, they had not been pre-treated with α-amylase so they included some accessible starch. Only 50%-70% of such flours is in the crystalline form that is resistant to mammalian α-amylase (23); the rest is “sensitive” or “accessible” to host enzymes. As a negative control we therefore included a maize starch that is completely digestible by human α-amylase so it should be broken down in the small intestine and not affect colonic communities. It was added at a low dose (equivalent to the sensitive portion of the RMS) or a high dose (equivalent to all the glucose monomers in that resistant supplement). In neither case did this placebo produce any statistically significant changes in the microbiota composition or SCFA production during consumption. The third supplement was the fructose polymer, inulin, that is entirely resistant to digestion in the small intestine.

Consuming resistant RPS led to an increase in the average concentration of fecal butyrate (Table 1). Neither inulin nor RMS produced a significant change in butyrate. An important caveat is that the amount of amylose-resistant polysaccharide consumed was not equal across treatment groups. The inulin and Hi-Maize groups consumed approximately 20 and 20-24 grams, respectively, while the RPS group consumed approximately 28-34 grams. However based on preliminary data, the discrepancy is not sufficient to explain the lack of a butyrogenic effect from inulin and Hi-Maize. In a pilot study, we observed a significant increase in fecal butyrate in individuals consuming half the dose of RPS (24 g total, 14-17 g resistant; data not shown). The butyrogenic response to RPS appears due to the nature of the supplement, not just the amount of RS it contains. So the answer to our first research question is that the fiber supplements are not equally effective at stimulating levels of this health-promoting metabolite.

All the fermentable fiber supplements had some effect on the fecal community (Fig. 2). In individuals consuming RPS, the largest and most common change in V4 sequences was an increase in sequences attributed to *B. faecale/adolescentis/stercoris*. This contrasts with an earlier study of dietary supplementation with resistant starch Type 3, where *R. bromii* was the dominant responder (22, 36) and was therefore proposed to be a keystone species for the degradation of resistant starch (18). The different responses in our study could be due to differences in the supplements, as has been observed previously when Type 2 or Type 4 resistant starches are consumed (37). The higher abundance of bifidobacteria in our study is consistent with the known ability of *B. adolescentis* to degrade and ferment RPS *in vitro* (20), but the increase was not associated with a change in fecal butyrate (Fig. 3B), suggesting that these organisms are not effectively cross-feeding butyrate producers *in vivo* – even though they can *in vitro* (20, 28, 29). It is conceivable that *in vivo* production of butyrate from such cross-feeding requires additional time for interactions to be established in the gut microbiome. Longer-term studies are currently underway to assess this possibility. The microbiomes in a subset of individuals consuming RPS did respond with an increase in *R*. bromii-associated sequences. In this group there was an associated increase in fecal concentrations of butyrate so *R. bromii* may be considered a keystone degrader with regard to butyrate production in our cohort (Fig. 3B). The major fermentation products of *R. bromii* are acetate, H_2_ and CO_2_, but not butyrate (35) so there must be cross-feeding of a butyrogenic microbe. The most likely candidate is *Eubacterium rectale* because its abundance increased most consistently with that of *R. bromii* (Fig. 4). Two less-well characterized bacterial populations also increased in relative abundance in a few individuals consuming RPS. The increase in the *Clostridium* leptum-related Seq100 was not associated with an increase in butyrate (Fig. 3B), suggesting that it too may be a degrader but not a cross-feeder. The Seq176 increase was associated with a butyrate increase (Fig. 3B), which may be because it is both a degrader and a butyrate producer. The activity of its cultured relative *Clostridium chartatabidum* against RPS is not documented, but it is a rumen bacterium capable of degrading a variety of plant fibers and, it produces butyrate (33, 34). Attempts to culture the human strain are underway.

Individuals consuming RMS did not respond with an increase in the abundance of *B. faecale/adolescentis/stercoris* sequences. With this supplement, like previous studies (36), an increase in *R. bromii*-associated sequences was the most common response (Fig. 2). Unlike the RPS-induced *R. bromii* increase, the increase in *R. bromii* on RMS was not associated with a significant increase in fecal butyrate. The lack of a butyrogenic response to RMS was unexpected because the supplement has led to increased fecal butyrate in animal models, though over four weeks (12). We speculate that the crystal structures of the resistant starches from the two plants are different because the RPS is phosphorylated (once every 200 glycosyl residues (38)). They may therefore be degraded with different efficiency or by different strains. Consequently more time may be required to develop cross-feeding interactions that generate measurable differences in fecal butyrate.

Inulin increased the relative abundance of four species of *Bifidobacterium*, consistent with the widespread occurrence of this degradative capability within the genus (26). There were also increases in the abundance of the butyrate producers *Anaerostipes hadrus* and *E. rectale* (Fig. 2), but they did not result in increased fecal butyrate. The *A. hadrus* may be feeding on the lactate and acetate produced by the more abundant Bifidobacteria on this supplement (31, 32) or it may be able to metabolize inulin itself. We note that the low pKa of lactic acid (pKA = 3.86) produced by the Bifid shunt (30) could reduce fecal pH and inhibit butyrogenic microbes that are sensitive to the lower pH. Subsequent utilization of lactic acid would restore pH but may extend the time required to see an effect. Or it could alter the distribution of microbial populations in the GI tract such that the balance of butyrate consumption and excretion by the host is affected (39). It may also be that lactate produced by *Bifidobacterium* species was converted to fermentation products other than butyrate. Notably propionate increased by 27% on average with inulin consumption, though the change was not statistically significant due to high variability. Other than *Bifidobacterium spp*. and *A. hadrus*, there were no significant increases in any other sequences (Supplementary Table 3), surprisingly including *F. prausnitzii* (which has the metabolic capacity to ferment inulin to butyrate (27)).

Thus we observed that all the fermentable fiber supplements we tested increased the relative abundances of some members of the fecal microbiota. Most of the affected sequences are associated with known degraders of resistant polysaccharides or producers of butyrate but we did uncover two Firmicutes not previously associated with fiber supplements. The organisms responding depended both on the individual and the supplement (research question 2).

Together our observations on SCFA and community composition changes suggest that the working model for stimulating butyrate production with fiber supplements (Fig. 1) is an over-simplification, in that fiber degradation does not always lead to butyrate production. The requirement for fiber breakdown by specialized primary degraders appears to hold, with lack of an appropriate degrader preventing butyrate increase. However, not all degraders lead to a butyrogenic response. The lack of enhanced butyrate production in *Bifidobacterium-responding* microbiomes suggests that while bifidobacteria are particularly effective at using some fiber supplements, they do not establish cross-feeding reactions with butyrogenic populations as readily as Ruminococcus-responsive microbiomes. Furthermore, the concept of a wide variety of butyrate-producing organisms having ready access to the products of degradation and fermentation is not supported. Even though many of these organisms can be cross-fed by primary degraders *in vitro* (20, 22, 28, 29), the *in vivo* scenario seems much more restricted. The only butyrate producer whose abundance increased with a primary degrader (Fig. 4) and was associated with higher fecal butyrate (Fig. 5) is *E. rectale*. This organism has a multi-protein, cell-wall attached system for degrading sensitive starch (40). It includes carbohydrate binding domains that enable it to attach to RS granules (but not degrade them), as well as ABC transporter proteins that capture oligosaccharide degradation products of the same size as those preferred for growth of *R. bromii* (22). It has been shown to co-localize with *R. bromii* on starch granules (24) and to grow on the products of RS degradation by *R. bromii* in vitro (22). It thus has the characteristics of a preferred partner in converting RPS to butyrate (research question 3). A single organism, such as *C. chartatabidum*, may be an alternative route, but it occurs in fewer individuals. If the *C. chartatabidum-related* organism from this cohort is indeed a primary degrader of RPS that produces butyrate, it would be an appealing probiotic to give in combination with RPS to enhance butyrate production in a larger percentage of individuals.

**Figure 5.**
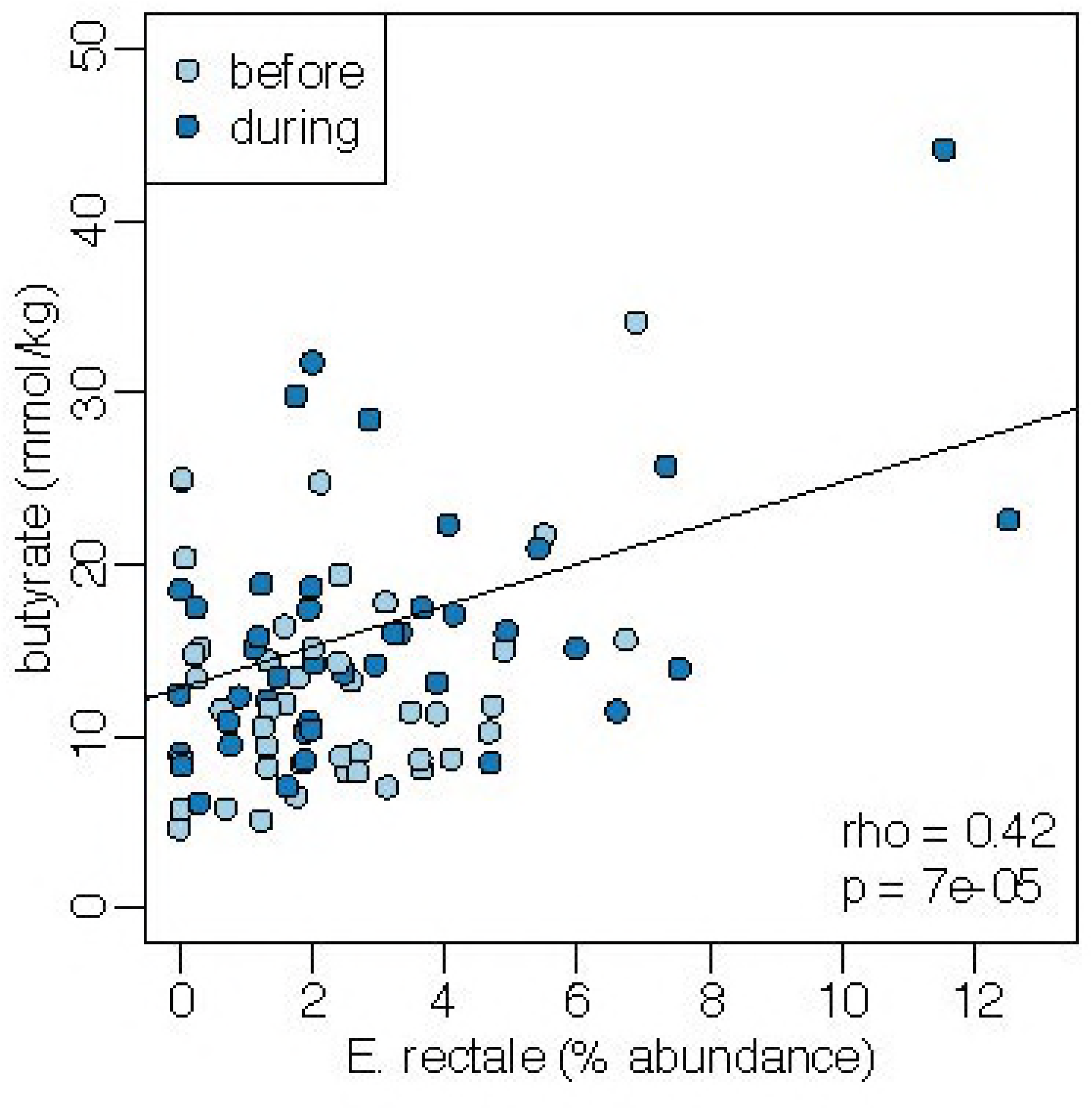
Positive relationship between fecal butyrate concentrations and the relative abundance of sequences characterized as *E. rectale* both before and during dietary supplementation with RPS.

To improve the efficacy of dietary supplements like these, it may be necessary to personalize them according to an individual’s gut microbiota (41). The presence of *R. bromii* or *C. chartatabidum* suggests whether a gut microbiome would yield increased butyrate concentrations following short-term (2-week) supplementation with RPS (research question 4). Individuals without *R. bromii* in their gut microbiome may benefit from a probiotic supplementation with *R. bromii* to increase the likelihood of a butyrogenic response to RPS. There may also be a synergistic effect of combining RPS with both *R. bromii* and *E. rectale* to maximize the butyrogenic effect of the supplement. In contrast, microbiomes with high levels of bifidobacteria are less likely to increase butyrate production in response to RPS (or inulin), at least in the short term. In these microbiomes, a different supplement or combination of supplements may be needed, or a longer period of time may be required for the microbiome to respond to the supplement. Such considerations are necessary when attempting to effect a particular change in the highly variable structures of human gut communities.

## MATERIALS AND METHODS

### Study Participants

Study participants were recruited through the Authentic Research Sections of the introductory biology laboratory course at the University of Michigan (BIO173). Individuals with self-reported history of inflammatory bowel syndrome, inflammatory bowel disease, or colorectal cancer were excluded from the study, as were individuals who had taken antibiotics within the last 6 months. Pre- or probiotic usage was not an exclusion criterion for the study. Nor was the amount of fiber already being consumed. All participants gave written, informed consent prior to participating in the study. Participants under the age of 18 were granted permission by a parent or legal guardian. Participants ranged in age from 17 to 29, with a median age of 19. This study was approved by the Institutional Review Board of the University of Michigan Medical School (HUM00094242 and HUM00118951) and was conducted in compliance with the Helsinki Declaration.

The participants were randomly assigned to study groups. During the first semester they were blinded to the supplement they were taking; in subsequent studies they were informed of its identity. Researchers analyzing samples were always blinded to the supplement associated with the samples. Some participants were excluded from the analysis because fewer than three samples were collected or successfully analyzed either before or during dietary supplementation.

### Study Design

This replicated intervention study was conducted during four separate academic semesters from the fall of 2015 to the spring of 2017. It was a parallel design with different but similar groups taking experimental or control supplements. Replicated baseline data were collected for each individual during the first week of each study. Each participant collected 3 to 4 fecal samples on separate days during this period. During the second week, participants underwent a 4-to 7-day transition phase, that began with consumption of a half dose of the supplement before taking the full dose. No fecal samples were collected during the transition phase. In the third week of the study participants continued taking the full dose of their assigned supplement until they had collected 3 to 4 fecal samples on separate days.

### Dietary Supplements

Four different supplements were tested in this study; resistant potato starch (RPS, Bob’s Red Mill, Milwaukee OR), Hi-Maize 260 resistant corn starch (RMS, manufactured by Ingredion Inc., Westchester, IL and distributed by myworldhut.com), inulin isolated from chicory root (Swanson Health Products, Fargo, ND), and amylase-accessible corn starch (Amioca powder, Skidmore Sales and Distribution, West Chester OH). Preliminary studies suggested that 40-48 g of supplement could be comfortably consumed per day. This amount was therefore used for the studies with RPS and RMS. However, it should be noted that these supplements do not contain the same proportion of amylase-resistant starch: RPS contains approximately 70% RS (Type 2) by weight and RMS approximately 50% (23). Subjects therefore consumed 28 – 34 grams of RPS or 20 – 24 grams of RMS per day. The inulin supplement, which is resistant to human enzymes (42), was consumed at 20 g/day to provide a similar amount of resistant polysaccharide. The accessible starch supplement was given at either 40 g/day (n=15) to provide a similar amount of total carbohydrates or at 20 g/day (n=24) to provide approximately the same amount of host-accessible carbohydrates. Consumption of the supplements was split into two half doses per day and logged on the study website. Participants were provided with a shaker bottle to aid in mixing the supplement with water, however they were permitted to consume the supplement with any type of food or beverage. Participants were instructed not to add the supplement to any warm food or beverage. Aside from the supplement, participants maintained their normal diet throughout the study.

### Fecal Collection

Fecal samples were collected by participants as described previously (10). Participants collected approximately half a gram of fecal material into an OMNIgene-Gut^®^ (DNA Genotek) collection kit, following manufacturer instructions. Collection tubes were transferred to a −20°C freezer within 24 hours of collection and stored at −20°C until thawed for DNA and metabolite extractions. Collection tubes were weighed before and after fecal collection, to determine the weight of the fecal material collected.

### SCFA Quantification

To extract SCFAs, 1 ml of fecal suspension was transferred into a 2 ml 96-well V-bottom collection plate and centrifuged at 4,500 x G for 15 minutes at 4°C. 200 μl of supernatant fractions were successively filtered through 1.20, 0.65, and 0.22 μm 96-well filter plates at 4°C. Filtrates were transferred into 1.5 ml screw cap vials containing 100 μl inserts in preparation for analysis by high-performance liquid chromatography (HPLC). Samples were analyzed in a randomized order. Quantification of SCFAs was performed using a Shimadzu HPLC system (Shimadzu Scientific Instruments, Columbia, MD) that included an LC-10AD vp pump A, LC-10AD vp pump B, degasser DGU-14A, CBM-20A, autosampler SIL-10AD HT, Column heater CTO-10A(C) vp, UV detector SPD-10A(V) vp, and an Aminex HPX-87H column (Bio-Rad Laboratories, Hercules, CA). We used a mobile phase of 0.01 N H2SO4 at a total flow rate of 0.6 ml per minute with the column oven temperature at 50°C. The sample injection volume was 10 μl and each sample eluted for 40 minutes. Concentrations were calculated using a cocktail of short chain organic acids standards at concentrations of 20, 10, 5, 2.5, 1, 0.5, 0.25, and 0.1 mM. After correcting the baseline of chromatographs, the quality of peaks was assessed using peak width, relative retention time, and 5% width. Peaks that fell outside predetermined cutoffs for relative retention time and peak width were removed. Concentrations in experimental samples were normalized to the wet weight of fecal material.

### 16S rRNA gene sequencing

DNA was extracted from 250 μl of fecal suspension using the 96-well MagAttract PowerMicrobiome DNA Isolation kit (Qiagen) and EpMotion liquid handling systems (Eppendorf). The V4 region of the bacterial 16S rRNA gene was amplified and sequenced as described previously using 2 x 250 base pair paired end kits on the Illumina MiSeq sequencing platform (43). Samples were assigned randomly to different runs each semester, with 8 separate DNA sequencing runs in total. Sequences were curated using the mothur software package as described previously (43, 44). Briefly, paired end reads were merged into contigs, screened for sequencing errors, and aligned to the SILVA bacterial SSU reference database. Aligned sequences were screened for chimeras and classified using the Ribosomal Database Project database. Sequences classified as mitochondria, chloroplasts, or archaea were removed. Sequences of interest were further identified using BLAST to align against the 16S rRNA gene sequences database. Unless stated otherwise, species designations indicate 100% identity to a single species in the database. The number of sequences per sample was rarified to 3000 to prevent biases from uneven sampling.

### Statistical analyses

All statistical analyses were performed using R (version 3.2.4) via RStudio (version 1.0.136). To determine whether there was a significant change in SCFA concentration from before to during fiber supplementation, we performed repeated measures ANOVAs on SCFA concentrations in all fecal samples from individuals consuming each supplement. For other analyses, we used the median SCFA concentration within each individual at each time point. Changes in the overall community structure in response to supplements were assessed using a within-subjects PERMANOVA on Bray-Curtis distances. Diversity was compared using repeated measures ANOVA on the inverse Simpson index for each sample. For other comparisons of the relative abundance of microbiota we used the average relative abundance within each individual at each time point, yielding a single average community structure for each individual before and during supplementation. Organisms that increased the most in response to each supplement were identified using one-tailed paired Wilcoxon-tests on the average abundances in each subject before and during supplementation with Benjamini-Hochberg correction for multiple comparisons (45). P-values in Figure 2 were corrected for the multiple tests applied to those 14 species, which were *a priori* expected to important for fiber degradation and/or butyrate production, while p-values in Supplemental Tables 1-4 were corrected for test across all 500 sequences. Correlation between the changes in abundance of primary degraders and butyrate producers (Fig. 4) were calculated using Spearman correlation with Benjamini-Hochberg correction for multiple comparisons. Raw sequencing reads and metadata, including SCFA concentrations, are available through the NCBI Short Read Archive under accession number SRP128128.

## ACKNOWLEDGEMENTS

This research was supported by grants from the Procter and Gamble Company, Howard Hughes Medical Institute (52008119), and the University of Michigan Host Microbiome Initiative. The authors declare that they have no competing interests.

The authors thank the students of Biology 173 who participated in this study and the University of Michigan Microbial Systems Molecular Biology Laboratory for performing 16S rRNA gene library preparation and sequencing.

